## Supplementary material for "A MademoiseLLE domain binding platform links the key RNA transporter to endosomes": Supplemenatary Table S1 to S9

$ shared first authorship

* Shared corresponding authorship

[1] Institute of Microbiology, Heinrich Heine University Düsseldorf, Cluster of Excellence on Plant Sciences, 40204 Düsseldorf, Germany

[2] John von Neumann Institute for Computing (NIC), Jülich Supercomputing Centre (JSC), Institute of Biological Information Processing (IBI-7: Structural Bioinformatics), and Institute of Bio- and Geosciences (IBG-4: Bioinformatics), Forschungszentrum Jülich GmbH, Wilhelm-Johnen-Str., 52425 Jülich, Germany

[3] Institute for Pharmaceutical and Medicinal Chemistry, Heinrich Heine University Düsseldorf, 40204 Düsseldorf, Germany

[4] Center for Structural Studies, Heinrich Heine University Düsseldorf, 40204 Düsseldorf, Germany

[5] Institute of Biochemistry I, Heinrich Heine University Düsseldorf, 40204 Düsseldorf, Germany

**Table of content:**

Appendix Table S1 Accession number of protein sequences……………...…………………02
Appendix Table S2 Data collection and refinement statistics………………...……………... 03
Appendix Table S3 Overall SAXS Data ……….……………………………………….......04
Appendix Table S4 Description of *U. maydis* strains used in this study………….…..……….05
Appendix Table S5 Generation of *U. maydis* strains used in this study ………………...…..06
Appendix Table S6 Description of plasmids used for *U. maydis* strain generation…...……. 09
Appendix Table S7 Description of plasmids used for recombinant expression in *E. coli*……10
Appendix Table S8 Description of plasmids used for yeast two-hybrid analyses…………....11
Appendix Table S9 DNA oligonucleotides used for plasmid generation……...……………..12

#### Table S1. Accession numbers for protein sequences used in multiple sequence alignment of MLLE domains

| Organism Name | Protein Name | Domain Name | Uniprot KB Number | Sequence coverage |
|---|---|---|---|---|
| *Homo sapiens* | Poly[A] binding protein, PABP | $MLLE^{PABP}$ | P11940 | 554 - 617 |
| *Triticum aestivum* | Poly[A] binding protein, PABP | $MLLE^{PABP}$ | P93616 | 564 - 627 |
| *Trypanosoma cruzi* | Poly[A] binding protein, PABP | $MLLE^{PABP}$ | Q27335 | 484 - 547 |
| *Leishmania major* | Poly[A] binding protein, PABP | $MLLE^{PABP}$ | E9AFX7 | 494 - 557 |
| *Saccharomyces cerevisiae* | Poly[A] binding protein, PABP | $MLLE^{PABP}$ | P04147 | 501 - 566 |
| *Homo sapiens* | E3 ubiquitin-protein ligase UBR5, EDD | $MLLE^{Ubr5}$ | O95071 | 2390 - 2452 |
| *Rattus norvegicus* | E3 ubiquitin-protein ligase UBR5 | $MLLE^{Ubr5}$ | Q62671 | 2380-2442 |
| *Ustilago maydis* | Poly[A] binding protein, Pab1 | $MLLE^{Pab1}$ | Q4P8R9 | 567 - 630 |
| *Ustilago maydis* | Rrm4 | $MLLE3^{Rrm4}$ | A0A0D1DWZ5 | 727 - 792 |
| *Ustilago maydis* | Rrm4 | $MLLE2^{Rrm4}$ | A0A0D1DWZ5 | 564 - 629 |
| *Ustilago maydis* | Rrm4 | $MLLE1^{Rrm4}$ | A0A0D1DWZ5 | 462 – 528 |

### Table S2. Data collection and refinement statistics

| | $\mathbf{MLLE2^{Rrm4}}$ |
|---|---|
| Wavelength | 0.979340 |
| Resolution range | 33.51 - 2.6 (2.693 - 2.6) |
| Space group | P 43 21 2 |
| Unit cell | 53.455 53.455 144.873 90 90 90 |
| Total reflections | 50419 (4828) |
| Unique reflections | 6898 (665) |
| Multiplicity | 7.3 (7.3) |
| Completeness (%) | 98.34 (98.08) |
| Mean I/sigma(I) | 17.90 (2.79) |
| Wilson B-factor | 70.38 |
| R-merge | 0.08252 (0.9043) |
| R-meas | 0.08902 (0.9692) |
| R-pim | 0.03265 (0.3441) |
| CC1/2 | 0.997 (0.859) |
| CC* | 0.999 (0.961) |
| Reflections used in refinement | 6879 (665) |
| Reflections used for R-free | 688 (65) |
| R-work | 0.2189 (0.3054) |
| R-free | 0.2646 (0.3718) |
| CC(work) | 0.969 (0.844) |
| CC(free) | 0.970 (0.761) |
| Number of non-hydrogen atoms | 1003 |
| Macromolecules | 1002 |
| Solvent | 1 |
| Protein residues | 131 |
| RMS(bonds) | 0.010 |
| RMS(angles) | 1.24 |
| Ramachandran favored (%) | 96.85 |
| Ramachandran allowed (%) | 2.36 |
| Ramachandran outliers (%) | 0.79 |
| Rotamer outliers (%) | 1.79 |
| Clashscore | 5.87 |
| Average B-factor | 76.62 |
| Macromolecules | 76.62 |
| Solvent | 78.27 |

Statistics for the highest-resolution shell are shown in parentheses.

### Table S3: Overall SAXS Data

| SAXS Device | BM29, ESRF Grenoble [1, 2] | |
|---|---|---|
| **Data collection parameters** | | |
| Detector | PILATUS 2 M | |
| Detector distance (m) | 2.827 | |
| Beam size | 200 µm x 200 µm | |
| Wavelength (nm) | 0.099 | |
| Sample environment | Quartz capillary,1 mm ø | |
| s range (nm-1)‡ | 0.025–6.0 | |
| Exposure time per frame (s) | 1 (10 frames each concentration) | |
| **Sample** | **H-Rrm4-NT4** | **G-Rrm4** |
| Organism | *Ustilago maydis* | *Ustilago maydis* |
| UniProt ID and range | A0A0D1DWZ5 | A0A0D1DWZ5 |
| Mode of measurement | Batch | Batch |
| Temperature (°C) | 10 | 10 |
| Protein buffer | 20mM Hepes pH 8.0, 200mM NaCl, 1mM βME | 20mM Hepes pH 8.0, 200mM NaCl, 1mM βME |
| Protein concentration (mg/ml) | 0.6 | 0.6 |
| **Structural parameters** | | |
| I(0) from P(r) | 41.90 | 103.50 |
| Rg (real-space from P(r)) (nm) | 5.55 | 8.99 |
| I(0) from Guinier fit | 43.34 | 104.01 |
| s-range for Guinier fit (nm-1) | 0.060 – 0.230 | 0.054 – 0.147 |
| Rg (from Guinier fit) (nm) | 5.60 | 8.78 |
| Points from Guinier fit | 4 - 37 | 3 - 21 |
| Dmax (nm) | 18.49 | 30.74 |
| POROD volume estimate (nm3) | 122.73 | 586.72 |
| **Molecular mass (kDa)** | | |
| From I(0) | 43.34 | 104.01 |
| From MoW2 [3] | 22.14 | 74.22 |
| From Vc [4] | 36.57 | 147.88 |
| From POROD | 61.37 – 76.71 | 293.36 – 366.70 |
| From sequence | 40.33 | 110.95 |
| **Structure Evaluation** | | |
| EOM fit χ2 | 1.262 | 1.289 |
| Ambimeter score | 2.307 | 2.530 |
| **Software** | | |
| ATSAS Software Version [5] | 3.0.3 | |
| Primary data reduction | PRIMUS [6] | |
| Data processing | GNOM [7] | |
| Ensemble modelling | EOM [8] | |
| Structure evaluation | AMBIMETER [9] | |
| Model visualization | PyMOL [10] | |

‡$s = 4\pi\sin(\theta)/\lambda$, $2\theta$ – scattering angle, $\lambda$ – X ray-wavelength


1. Pernot, P., et al., *New beamline dedicated to solution scattering from biological macromolecules at the ESRF.* Journal of Physics: Conference Series, 2010. **247**(1): p. 012009.
2. Pernot, P., et al., *Upgraded ESRF BM29 beamline for SAXS on macromolecules in solution.* J Synchrotron Radiat, 2013. **20**(Pt 4): p. 660-4.
3. Fischer, H., et al., *Determination of the molecular weight of proteins in solution from a single small-angle X-ray scattering measurement on a relative scale.* Journal of Applied Crystallography, 2010. **43**: p. 101-109.
4. Rambo, R.P. and J.A. Tainer, *Accurate assessment of mass, models and resolution by small-angle scattering.* Nature, 2013. **496**(7446): p. 477-81.
5. Manalastas-Cantos, K., et al., *ATSAS 3.0: expanded functionality and new tools for small-angle scattering data analysis.* Journal of Applied Crystallography, 2021. **54**(1).
6. Konarev, P.V., et al., *PRIMUS: a Windows PC-based system for small-angle scattering data analysis.* Journal of Applied Crystallography, 2003. **36**: p. 1277-1282.
7. Svergun, D.I., *Determination of the Regularization Parameter in Indirect-Transform Methods Using Perceptual Criteria.* Journal of Applied Crystallography, 1992. **25**: p. 495-503.
8. Tria, G., et al., *Advanced ensemble modelling of flexible macromolecules using X-ray solution scattering.* IUCrJ, 2015. **2**(Pt 2): p. 207-17.
9. Petoukhov, M.V. and D.I. Svergun, *Ambiguity assessment of small-angle scattering curves from monodisperse systems.* Acta Crystallogr D Biol Crystallogr, 2015. **71**(Pt 5): p. 1051-8.
10. PyMOL, *The PyMOL Molecular Graphics System, Version 2.0 Schrödinger, LLC.* 2015.

#### Table S4: Description of *U. maydis* strains used in this study

| Strain name with code | Locus | Progenitor strain | Short description |
|---|---|---|---|
| AB33 (UMa133) | *b* | FB2 | *Pnar:bW2bE1*, expression of active b heterodimer under control of the *nar1* promoter, strain grows filamentous upon changing the nitrogen source. |
| AB33rrm4Δ/upa1-gfp (UMa2769) | *rrm4* *upa1* | AB33rrm4-Cherry/ upa1-gfp | carrying a deletion of *rrm4* and expressing Upa1 C-terminally fused to eGfp |
| AB33upa1-gfp/rrm4-kat (UMa2976) | *rrm4* *upa1* | AB33rrm4Δ/upa1-gfp | expressing Upa1 C-terminally fused to eGfp and Rrm4 C-terminally fused to mKate2 |
| AB33upa1-gfp/rrm4-m1Δ-kat (UMa2977) | *rrm4* *upa1* | AB33rrm4Δ/upa1-gfp | expressing Upa1 C-terminally fused to eGfp and Rrm4-M1Δ-C-terminally fused to mKate2. Like rm4-kat but carrying the deletion of 1st MLLE domain. Residues of Rrm4 from 447 to 540 were replaced with a HAtag-HRV3C protease recognition site. |
| AB33upa1-gfp/rrm4-m2Δ-kat (UMa2978) | *rrm4* *upa1* | AB33rrm4Δ/upa1-gfp | expressing Upa1 C-terminally fused to eGfp and Rrm4-M2Δ-C-terminally fused to mKate2. Like rrm4-kat but carrying the deletion of 2nd MLLE domain. Residues of Rrm4 from 547 to 644 were replaced with a HAtag-HRV3C protease recognition site. |
| AB33upa1-gfp/rrm4-m3Δ-kat (UMa2979) | *rrm4* *upa1* | AB33rrm4Δ/upa1-gfp | expressing Upa1 C-terminally fused to eGfp and Rrm4-M3Δ-C-terminally fused to mKate2. Like rrm4-kat but carrying the deletion of 3rd MLLE domain. Residues of Rrm4 from 689 to 792 were replaced with a HAtag-HRV3C protease recognition site. |
| AB33upa1-gfp/rrm4-m1,2Δ-kat (Uma2981) | *rrm4* *upa1* | AB33rrm4Δ/upa1-gfp | expressing Upa1 C-terminally fused to eGfp and Rrm4-M1,2Δ-C-terminally fused to mKate2. Like rrm4-kat but carrying the deletion of 1st and 2nd MLLE domains. Residues of Rrm4 from 447 to 644 were replaced with a HAtag-HRV3C protease recognition site. |
| AB33upa1-pl1m-gfp/rrm4-m1,2Δ-kat (Uma2982) | *rrm4* *upa1* | AB33rrm4Δ/upa1-pl1m-gfp | expressing Upa1-PL1m- C-terminally fused to eGfp, carries block mutations leading to the amino acid substitutions AASAAATAAS from residues 242-251 in the N-terminal PAM2L-motif (PAM2L-1). Rrm4-M1,2Δ C-terminally fused to mKate2 and carrying the deletion of 1st and 2nd MLLE domains. Residues of Rrm4 from 447 to 644 were replaced with a HAtag-HRV3C protease recognition site. |
| AB33upa1-pl2m-gfp/rrm4-m1,2Δ-kat (Uma2983) | *rrm4* *upa1* | AB33rrm4Δ/upa1-pl2m-gfp | expressing Upa1-PL2m- C-terminally fused to eGfp, carries block mutations leading to the amino acid substitutions AASAAATAAS from residues 949-958 in the C-terminal PAM2L-motif (PAM2L-2). Rrm4-M1,2Δ C-terminally fused to mKate2 and carrying the deletion of 1st and 2nd MLLE domains. Residues of Rrm4 from 447 to 644 were replaced with a HAtag-HRV3C protease recognition site. |
| AB33upa1-pl1,2m-gfp/rrm4-m1,2Δ-kat (Uma3177) | *rrm4* *upa1* | AB33rrm4Δ/upa1-pl1,2m-gfp | expressing Upa1-PL1,2m C-terminally fused to eGfp, carries block mutations leading to the amino acid substitutions AASAAATAAS from residues 242-251 in the N-terminal PAM2L-motif (PAM2L-1) and from residues 949-958 in the C-terminal PAM2L-motif (PAM2L-2). Rrm4-M1,2Δ C-terminally fused to mKate2 and carrying the deletion of 1st and 2nd MLLE domains. Residues of Rrm4 from 447 to 644 were replaced with a HAtag-HRV3C protease recognition site. |

| | | | |
|---|---|---|---|
| AB33upa1Δ/rrm4-kat<br>(Uma3179) | *rrm4*<br>*upa1* | AB33upa1-gfp/rrm4-kat | Carrying a deletion of *upa1* and Rrm4 C-terminally fused to mKate2 |
| AB33upa1-pl1,2m-gfp/rrm4-kat<br>(Uma3355) | *rrm4*<br>*upa1* | AB33rrm4Δ/upa1-pl1,2m-gfp | expressing Upa1-PL1,2m C-terminally fused to eGfp, carries block mutations leading to the amino acid substitutions AASAAATAAS from residues 242-251 in the N-terminal PAM2L-motif (PAM2L-1) and from residues 949-958 in the C-terminal PAM2L-motif (PAM2L-2). Rrm4 C-terminally fused to mKate2. |
| AB33upa1-pl1m-gfp/rrm4-kat<br>(UL46) | *rrm4*<br>*upa1* | AB33rrm4Δ/upa1-pl1m-gfp | expressing Upa1-PL1m C-terminally fused to eGfp, carries block mutations leading to the amino acid substitutions AASAAATAAS from residues 242-251 in the N-terminal PAM2L-motif (PAM2L-1) and Rrm4 C-terminally fused to mKate2. |
| AB33upa1-pl2m-gfp/rrm4-kat<br>(UL47) | *rrm4*<br>*upa1* | AB33rrm4Δ/upa1-pl2m-gfp | expressing Upa1-PL2m C-terminally fused to eGfp, carries block mutations leading to the amino acid substitutions AASAAATAAS from residues 949-958 in the C-terminal PAM2L-motif (PAM2L-2) and Rrm4 C-terminally fused to mKate2. |
| AB33upa1-pl2m-gfp/rrm4-m1Δ-kat<br>(UL48) | *rrm4*<br>*upa1* | AB33rrm4Δ/upa1-pl1,2m-gfp | expressing Upa1-PL2m C-terminally fused to eGfp, carries block mutations leading to the amino acid substitutions AASAAATAAS from residues 949-958 in the C-terminal PAM2L-motif (PAM2L-2). Rrm4-M1Δ C-terminally fused to mKate2. Like rrm4-kat but carrying the deletion of 1st MLLE domain. Residues of Rrm4 from 447 to 540 were replaced with a HAtag-HRV3C protease recognition site. |
| AB33upa1-pl2m-gfp/rrm4-m1,2Δ-kat<br>(UL49) | *rrm4*<br>*upa1* | AB33rrm4Δ/upa1-pl2m-gfp | expressing Upa1-PL2m C-terminally fused to eGfp, carries block mutations leading to the amino acid substitutions AASAAATAAS from residues 949-958 in the C-terminal PAM2L-motif (PAM2L-2). Rrm4-M1,2Δ C-terminally fused to mKate2 and carrying the deletion of 1st and 2nd MLLE domains. Residues of Rrm4 from 447 to 644 were replaced with a HAtag-HRV3C protease recognition site. |

#### Table S5: Generation of *U. maydis* strains used in this study

| Strains | Relevant genotype | Strain code | Reference | Transformed plasmid | Locus | Progenitor |
|---|---|---|---|---|---|---|
| AB33 | *a2 $P_{nar}$:bW2 bE1* | UMa 133 | Brachmann, 2001 | pAB33 | *b* | FB2 |
| AB33rrm4Δ/upa1-gfp | *rrm4 upa1-gfp* | UMa 2769 | this study | pRrm4Δ_genitR (pUMa1755) | *rrm4* | AB33rrm4-mCherry/upa1-gfp (UMa1594) |
| AB33upa1-gfp/rrm4-kat | *upa1-gfp rrm4-kat* | Uma 2976 | this study | pRrm4-kat-hygR (pUMa3908) | *rrm4* | AB33rrm4Δ/upa1-gfp (UMa2769) |
| AB33upa1-gfp/rrm4-m1Δ-kat | *upa1-gfp rrm4*-m1Δ-kat | UMa 2977 | this study | pRrm4-m1Δ-kat-hygR (pUMa4433) | *rrm4* | AB33rrm4Δ/upa1-gfp (UMa2769) |
| AB33upa1-gfp/rrm4-m2Δ-kat | *upa1-gfp rrm4-m2Δ-kat* | UMa 2978 | this study | pRrm4- m2Δ-kat-hygR (pUMa4434) | *rrm4* | AB33rrm4Δ/upa1-gfp (UMa2769) |
| AB33upa1-gfp/rrm4-m3Δ-kat | *upa1-gfp rrm4-m3Δ-kat* | UMa 2979 | this study | pRrm4-m3Δ-kat-hygR (pUMa4435) | *rrm4* | AB33rrm4Δ/upa1-gfp (UMa2769) |
| AB33upa1-gfp/rrm4-m1,2Δ-kat | *upa1-gfp rrm4-m1,2Δ-kat* | UMa 2981 | this study | pRrm4-m1,2Δ-kat-hygR (pUMa4578) | *rrm4* | AB33rrm4Δ/upa1-gfp (UMa2769) |
| AB33upa1-pl1m-gfp/ rrm4- m1,2Δ-kat | *upa1-pl1m-gfp rrm4-m1,2Δ-kat* | UMa 2982 | this study | pRrm4-m1,2Δ-kat-hygR (pUMa4578) | *rrm4* | AB33rrm4Δ/upa1-pl1m-gfp (UMa2766) |
| AB33upa1-pl2m-gfp/ rrm4-m1,2Δ-kat | *upa1-pl2m-gfp rrm4-m1,2Δ-kat* | UMa 2983 | this study | pRrm4-m1,2Δ-kat-hygR (pUMa4578) | *rrm4* | AB33rrm4Δ/upa1-pl2m-gfp (UMa2767) |
| AB33upa1-pl1,2m-gfp/ rrm4-m1,2Δ-kat | *upa1-pl1,2m-gfprrm4-m1,2Δ-kat* | UMa 3177 | this study | pRrm4-m1,2Δ-kat-hygR (pUMa4578) | *rrm4* | AB33rrm4Δ/upa1-pl1,2m-gfp (UMa2768) |
| AB33upa1Δ/rrm4-kat | *upa1Δ rrm4-kat* | UMa 3179 | this study | pUpa1Δ-genitR (pUMa1915) | *upa1* | AB33upa1-gfp/rrm4-kat (UMa2976) |

| | | | | | | |
|---|---|---|---|---|---|---|
| AB33upa1-pl1,2m-gfp/ rrm4-kat | *upa1-pl1,2m-gfp* *rrm4-kat* | UMa 3355 | this study | pRrm4-kat-hygR (pUMa3908) | *rrm4* | AB33rrm4Δ/upa1-pl1,2m-gfp (UMa2768) |
| AB33upa1-pl1m- gfp/ rrm4-kat | *upa1-pl1m-gfp* *rrm4-kat* | UL46 | this study | pRrm4-kat-hygR (pUMa3908) | *rrm4* | AB33rrm4Δ/upa1-pl1m-gfp (UMa2766) |
| AB33upa1-pl2m-gfp/ rrm4-kat | *upa1-pl2m-gfprrm4-kat* | UL47 | this study | pRrm4-kat-hygR (pUMa3908) | *rrm4* | AB33rrm4Δ/upa1-pl2m-gfp (UMa2767) |
| AB33upa1-pl2m-gfp/ rrm4-m1Δ-kat | *upa1-pl2m-gfp* *rrm4-m1Δ-kat* | UL48 | this study | pRrm4-m1Δ-kat-hygR (pUMa4433) | *rrm4* | AB33rrm4Δ/upa1-pl2m-gfp (UMa2767) |
| AB33upa1-pl2m-gfp/ rrm4-m2Δ-kat | *upa1-pl2m-gfp* *rrm4-m2Δ-kat* | UL49 | this study | pRrm4-m2Δ-kat-hygR (pUMa4434) | *rrm4* | AB33rrm4Δ/upa1-pl2m-gfp (UMa2767) |

#### Table S6: Description of plasmids used for *U. maydis* strain generation

| Plasmid | pUMa | Resistance cassette | Short description |
|---|---|---|---|
| pRrm4Δ | 1755 | genitR (G418 resistance - SfiI insert of pMF1g) Baumann et al., 2012 | Plasmid vector for generating deletion mutants of *rrm4*. |
| pUpa1Δ_genitR | 1915 | genitR | Plasmid vector for generating deletion mutants of *upa1*. (Pohlmann et al., 2015) |
| pRrm4-kat-hygR | 3908 | hygR (Hygromycin resistance - SfiI insert of pMF1h) Brachmann et al., 2004 | Plasmid vector for the expression of Rrm4 C-terminally fused to mKate2. The mKate2 cassette contains the Tnos terminator and the Hyg resistance. The entire coding sequence for the fusion protein is flanked by a 1025 bp upstream region and a 1396 bp downstream region for homologous recombination. |
| pRrm4-m1Δ-kat-hygR | 4433 | hygR | Plasmid vector for the expression of Rrm4-M1Δ C-terminally fused to mKate2. Like pRrm4-mK-HygR, but carrying the deletion of 1st MLLE domain. Residues of Rrm4 from 447 to 540 were replaced with a HAtag-HRV3C protease recognition site. |
| pRrm4-m2Δ-kat-hygR | 4434 | hygR | Plasmid vector for the expression of Rrm4-M2Δ C-terminally fused to mKate2. Like pRrm4-mK-HygR, but carrying the deletion of the 2nd MLLE domain. Residues of Rrm4 from 547 to 644 were replaced with a HAtag-HRV3C protease recognition site. |
| pRrm4-m3Δ-kat-hygR | 4435 | hygR | Plasmid vector for the expression of Rrm4-M3Δ C-terminally fused to mKate2. Like pRrm4-mK-HygR, but carrying the deletion of the 3rd MLLE domain. Residues of Rrm4 from 689-792 were replaced with a HAtag-HRV3C protease recognition site. |
| pRrm4-m1,2Δ-kat-hygR | 4578 | hygR | Plasmid vector for the expression of Rrm4-M1,2Δ C-terminally fused to mKate2. Like pRrm4-mK-HygR, but carrying the deletion of 1st and 2nd MLLE domains. Residues of Rrm4 from 447 to 644 were replaced with a HAtag-HRV3C protease recognition site. |

#### Table S7: Description of plasmids used for recombinant expression in *E. coli*

| Plasmid | pUMa | Short description |
|---|---|---|
| pGEX-G-Pab1-MLLE | 2187 | Plasmid for the expression of the G-Pab1-MLLE. C terminal region of Pab1 comprising amino acid residues 566-651 were N-terminally fused to a GST-tag. (Pohlmann et al., eLife 2015) |
| pGEX-G-Rrm4-NT4 | 3920 | Plasmid for the expression of the G-Rrm4-NT4. C terminal region of Rrm4 comprising amino acid residues 421 to 792 was N-terminally fused to a GST-tag. (Pohlmann et al., eLife 2015) |
| pGEX-G-Rrm4-NT4-M1Δ | 4616 | Plasmid for the expression of the G-Rrm4-NT4-M1Δ. Same as pUMa3920 but carrying the deletion of 1st MLLE domain. Residues of Rrm4 from 447 to 540 were replaced with a HAtag-HRV3C protease recognition site. |
| pGEX-G-Rrm4-NT4-M2Δ | 4617 | Plasmid for the expression of the G-Rrm4-NT4-M2Δ. Same as pUMa3920 but carrying the deletion of the 2nd MLLE domain. Residues of Rrm4 from 547 to 644 were replaced with a HAtag-HRV3C protease recognition site. |
| pGEX-G-Rrm4-NT4-M3Δ | 4618 | Plasmid for the expression of the G-Rrm4-NT4-M3Δ. Same as pUMa3920 but carrying the deletion of the 3rd MLLE domain. Residues of Rrm4 from 689 to 792 were replaced with a HAtag-HRV3C protease recognition site. |
| pGEX-G-Rrm4-NT4-M1,2Δ | 4619 | Plasmid for the expression of the G-Rrm4-NT4-M1,2Δ. Same as pUMa3920 but carrying the deletion of 1st to 2nd MLLE domains. Residues of Rrm4 from 447 to 644 were replaced with a HAtag-HRV3C protease recognition site. |
| pET28-HS-PAM2$^{Upa1}$ | 4296 | Plasmid for the expression of the PAM2 motif of Upa1 (SQSTLSPNASVFKPSRS) as a fusion protein with an N terminal 6xHis-Sumo-tag. |
| pET28-HS_PAM2L1$^{Upa1}$ | 4297 | Plasmid for the expression of PAM2L1 motif of Upa1 (EAADQEEDQDDFVYPGAD) as a fusion protein with an N terminal 6xHis-Sumo-tag. |
| pET28-HS-PAM2L2$^{Upa1}$ | 4298 | Plasmid for the expression of PAM2L2 motif of Upa1 (DEDAADDDDDEFIYPNSY) as a fusion protein with an N terminal 6xHis-Sumo-tag. |
| pET22-H-Rrm4-NT4 | 3552 | Plasmid for the expression of H-Rrm4-NT4. C terminal region of Rrm4 comprising amino acid 421 to 792 were N-terminally fused to 6xHis-tag. |
| pGX-G-Rrm4 | 429 | Plasmid for the expression of G-Rrm4. Rrm4 full-length protein was N-terminally fused to GST. |

#### Table S8: Description of plasmids used for yeast two-hybrid analyses

| Plasmid | Plasmid code | Gene | Short description |
|---|---|---|---|
| pGADT7-DS | pUMa1624 | | Plasmid for the expression of hybrid proteins, N-terminally fused to a nuclear localisation signal (NLS) of the simian virus 40 (SV40), followed by the Gal4 activation domain (aa 768-881) and an HA-epitope for Western Blot detection. The resulting hybrid proteins are termed AD-“X”. For the positive selection of transformants on minimal medium, this plasmid carries a *LEU2* auxotrophy marker. This plasmid contains two diverse SfiI-restriction sites for cloning purposes (Dualsystems Biotech, Schlieren, Switzerland). |
| pGBKT7-SfiI MCS | pUMa1625 | | Plasmid for the expression of hybrid proteins, N-terminally fused to the Gal4 DNA-binding domain (aa 1-147), followed by a c-Myc-epitope for Western Blot detection. The resulting hybrid proteins are termed BD-“X”. For the positive selection of transformants on minimal medium, this plasmid carries a *TRP1* auxotrophy marker. This plasmid contains two diverse SfiI-restriction sites for cloning purposes. (Clontech Laboratories, Inc., Mountain View, CA, USA). |
| pGADT7-T | pUMa1636 | | Plasmid for the expression of an N-terminal AD-fusion of the large T-antigen of SV40. It interacts with BD-p53 as a positive control (Clontech). |
| pGBKT7-p53 | pUMa1638 | | Plasmid for the expression of an N-terminal BD-fusion of the murine p53. It interacts with AD-T as a positive control (Clontech). |
| pGBKT7-Lam | pUMa1637 | | Plasmid for the expression of an N-terminal BD-fusion with the human nuclear protein Lamin C, which shows no interaction with most proteins and serves as negative control (Clontech). |
| pGBKT7-Upa1-Gfp | pUL0128 | *upa1* | Plasmid for the expression of BD-Upa1-Gfp, where eGfp is fused C-terminally to the BD-Upa1-hybrid. |
| pGBKT7-Upa1-pl1m-Gfp | pUL0120 | *upa1* | Like GBKT7-Upa1-Gfp, expressing BD-Upa1-pl1-Gfp, where eGfp is fused C-terminally to the BD-Upa1-pl1m hybrid but carries block mutations leading to the amino acid substitutions AASAAATAAS from residues 242-251 in the N-terminal PAM2L-motif (PAM2L-1) of Upa1. |
| pGBKT7-Upa1-pl2m-Gfp | pUL0121 | *upa1* | Like pGBKT7-Upa1-Gfp, expressing BD-Upa1-pl2-Gfp, where eGfp is fused C-terminally to the BD-Upa1-pl2m hybrid. but carries block mutations leading to the amino acid substitutions AASAAATAAS from residues 949-958 in the C-terminal PAM2L-motif (PAM2L-2) of Upa1. |
| pGBKT7-Upa1-pl1,2m-Gfp | pUL0122 | *upa1* | Like GBKT7-Upa1-Gfp, expressing BD-Upa1-pl1,2-Gfp, where eGfp is fused C-terminally to the BD-Upa1-pl1,2m hybrid but carries block mutations leading to the amino acid substitutions AASAAATAAS from residues 242-251 in the N-terminal PAM2L-motif (PAM2L-1) and from residues 949-958 in the C-terminal PAM2L-motif (PAM2L-2). |
| pGADT7-Rrm4-kat | pUL0112 | *rrm4* | Plasmid for the expression of AD-Rrm4-kat, where mKate2 is fused C-terminally to the AD-Rrm4-hybrid. |
| pGADT7-Rrm4-m1Δ-kat | pUL0116 | *rrm4* | Plasmid for the expression of AD-Rrm4-M1Δ-kat where mKate2 is fused C-terminally to the AD-Rrm4-M1Δ hybrid. Like pGADT7-Rrm4-kat, but carrying the deletion of 1st MLLE domain. Residues of Rrm4 from 447 to 540 were replaced with a HAtag-HRV3 C protease recognition site. |
| pGADT7-Rrm4-m2Δ-kat | pUL0117 | *rrm4* | Plasmid for the expression of AD-Rrm4-M2Δ-kat, where mKate2 is fused C-terminally to the AD-Rrm4-M2Δ hybrid. Like pGADT7-Rrm4-kat, but carrying the deletion of 2nd MLLE domain. Residues of Rrm4 from 547 to 644 were replaced with a HAtag-HRV3C protease recognition site. |
| pGADT7-Rrm4-m1,2Δ-kat | pUL0118 | *rrm4* | Plasmid for the expression of AD-Rrm4-M1,2Δ-kat, where mKate2 is fused C-terminally to the AD-Rrm4-M1,2Δ hybrid. Like pGADT7-Rrm4-kat, but carrying the deletion of 1st and 2nd MLLE domains. Residues of Rrm4 from 447 to 644 were replaced with a HAtag-HRV3C protease recognition site. |
| pGADT7-Rrm4-m3Δ-kat | pUL0119 | *rrm4* | Plasmid for the expression of AD-Rrm4-M3Δ-kat, where mKate2 is fused C-terminally to the AD-Rrm4-M3Δ hybrid. Like pGADT7-Rrm4-kat, but carrying the deletion of 3rd MLLE domain. Residues of Rrm4 from 689-792 were replaced with a HAtag-HRV3C protease recognition site. |

#### Table S9: DNA oligonucleotides used in this study

| Designation | Nucleotide sequence (5' --> 3') | Remarks |
|---|---|---|
| oUM727 | GTATTCGAGCCAAGCATCTACGTATGTCGACCCTTGCAACC | Rrm4-internal-Gibson cloning-fwd |
| oAB354 | GGGCCCCTGGAACAGTACTTCCAGGGCGTAGTCGGGCACGTCGTAAGGGTAAGGCACACCTGCTTTGAAG | Rrm4-M1Δ_Gibson cloning-rev |
| oAB355 | TACCCTTACGACGTGCCCGACTACGCCCTGGAAGTACTGTTCCAGGGGCCCCTGTCTGCTGAACACCCAGC | Rrm4-M1Δ_Gibson cloning-fwd |
| oAB359 | CGATCGCCGGGCGGCCGGCGCGCCACCGGTTTAGCGGTGACCGAGTTTCGAGG | mKate2-rev |
| oAB345 | GGGCCCCTGGAACAGTACTTCCAGGGCGTAGTCGGGCACGTCGTAAGGGTATGCAGGAAGCGCAGCAAGCG | Rrm4-M3Δ_Gibson cloning-rev |
| oAB346 | TACCCTTACGACGTGCCCGACTACGCCCTGGAAGTACTGTTCCAGGGGCCCGCGGCCAACGCGGCCACCATGGTG | Rrm4-M3Δ_Gibson cloning-fwd |
| oAB356 | GGGCCCCTGGAACAGTACTTCCAGGGCGTAGTCGGGCACGTCGTAAGGGTATGGGTGTTCAGCAGACAGTG | Rrm4-M2Δ_Gibson cloning-rev |
| oAB357 | TACCCTTACGACGTGCCCGACTACGCCCTGGAAGTACTGTTCCAGGGGCCCAGCGCTCCGGTGCCATTGTC | Rrm4-M2Δ-Gibson cloning-fwd |
| oAB312 | CATGCCATGGCCAGCAGCAACAGTCCGCCCAC | NcoI_Rrm4-NT4-fwd |
| oAB45 | CGGCCATATGGGCAGCAGCCATCATC | pET28 vector-ORF-fwd |
| oAB46 | CTCACTCGAGTTAGGATCGGGACGGCTTGAAGACGGAGGCGTTGGGAGACAAGGTGCTTTGCGAACCACCAATCTGTTCTCTGTGAGC | Sumo-PAM2-XhoI-rev |
| oAB47 | CTCACTCGAGTTAGTCGGCTCCTGGGTAGACAAAGTCATCTTGATCTTCCTCTTGGTCTGCAGCCTCACCACCAATCTGTTCTCTGTGAG | Sumo-PAM2L1-XhoI-rev |
| oAB48 | CTCACTCGAGTTAGTACGAGTTCGGGTAGATGAATTCATCGTCATCGTCATCGGCCGCATCCTCGTCACCACCAATCTGTTCTCTGTGAG | Sumo-PAM2L2-XhoI-rev |
| oMB696 | TCGACTCGAGTCACTTGTTCAGACCTGCAGC | Rrm4- XhoI-rev |
| oUP605 | GGGAATTCCATATGCATCATCATCATCATCACAGCAGCAACAGTCCGCCCAC | NdeI-6xHis-Rrm4-NT4 fwd |
